# The human genome’s vocabulary as proposed by the DNA language model GROVER

**DOI:** 10.1101/2023.07.19.549677

**Authors:** Melissa Sanabria, Jonas Hirsch, Anna R. Poetsch

**Affiliations:** Biomedical Genomics, Biotechnology Center, Center for Molecular and Cellular Bioengineering, Technische Universität Dresden

## Abstract

Large Language Models (LLMs) on natural language have achieved a level of performance that allows the generation of coherent and syntactically correct text. DNA sequence of genomes follows rules similar to natural language, but a distinguishing factor is the absence of a concept analogous to words. We established byte-pair tokenization on the human genome and trained a foundation language model called GROVER (“<u>G</u>enome <u>R</u>ules <u>O</u>btained <u>V</u>ia <u>E</u>xtracted <u>R</u>epresentations”) to select the optimal vocabulary with a custom fine-tuning task of next-k-mer prediction. We thus defined a dictionary of words/tokens in the human genome that best carries the information content for DNA language models. Analyzing GROVER’s learned representations, we observed that token embeddings primarily encode information related to their frequency, sequence content, and length. Some tokens are almost exclusively localized in repeats, while the vast majority widely distributes over the genome. The model also learns context and lexical ambiguity. Average embeddings of genomic regions relate to functional genomics annotation and thus indicate that GROVER has learned these structures purely from the contextual relationships of tokens. That we can extract functional annotations from the genome, purely based on sequence representation to the trained model, highlights the extent of information content encoded by the sequence. This is supported by fine-tuning tasks on genome biology with questions of promoter identity and protein-DNA binding. GROVER learns sequence context, a sense for grammatical structures and language rules in the genome. This knowledge can be extracted and used to compose a grammar book for the code of life.

## Introduction

Genomes carry the instruction manuals for whole organisms. The first draft of the human genome has been available for >20 years ^1^, and genome sequences of multiple species have become available since. We know the letters and the text, but we still understand very little about the “genetic code”. DNA triplets encode for amino acids ^2^ in 1-2% of the genome ^1^, but there are additional layers to the “code of life”. There is encryption on how those genes are regulated, how transcripts and RNAs are structured, how they function and keep stable, how, where, and when the genome replicates, and also how all this is achieved with keeping the genome stable and functional. Extracting comprehensively these different layers of code requires complex algorithms that only recently became available through progress in natural language processing (NLP). Natural language can be studied with large language models (LLMs) that are based on transformer architectures ^3^ and very suitable for tasks involving text data due to their unprecedented performance and transparency. Pretrained LLMs like GPT-3 ^4^ and successors can also function as foundation models that can be fine-tuned with different classification, regression, or generative tasks. These models have changed how we can view language and they can be very useful for a variety of purposes. For example, they can make up very believable text, when given a generative task.

While genomes and DNA are very analogous to language through sequence structure that resembles grammar, syntax, and semantics, they also differ. First, there is no clearly defined direction in DNA sequence, unless viewed relative to biological processes like transcription or replication. Second, there is no natural definition of words. We know of motifs, such as sequence with preferences to be bound by specific transcription factors, or the triplet code that encodes for protein. However, when looking at the genome as a whole, there is no straight forward concept of words. To overcome those challenges for training transformer models on DNA, so called DNA language models, there have been several approaches. Some are aimed to address specific tasks, such as the modelling of gene expression with Enformer ^5^, a model that combines convolutional layers with transformer blocks. Through the convolutional layer, there is no definition of words necessary. Foundation models however are trained not directly on a specific genome biology task, but are first pretrained on masked token prediction and subsequently fine-tuned. This strategy requires the definition of discrete tokens, i.e. to build “words” from DNA. Available models include DNABERT ^6^ and Nucleotide transformer ^7^ that use a Bidirectional Encoder Representations from Transformers (BERT) ^8^ architecture and apply different strategies of generating the vocabulary. Nucleotide transformer uses as vocabulary 6mers and single nucleotides on edges of known sequence. DNABERT on the other hand uses k-mers of 3, 4, 5, and 6 nucleotides in length for four different models, of which the 6mer model performs best ^6^. The k-mers have overlaps and the training is designed for the central nucleotide of a masked sequence not to overlap with any unmasked tokens. As a consequence, the model largely learns the sequence of the tokens, rather than larger sequence context ^9^.

How to choose the right vocabulary for training a DNA language model has been challenging. On the one hand, the tokens should have an appropriate length so that they capture the language structure of the genome. However, if this length is chosen as a constant, frequencies of tokens become very heterogeneous. For 6mers, for example, the frequency of each token ranges from ca. 10^4^ to 10^7^ occurrences in the human genome (hg19). Such a frequency imbalance can inhibit model training through Rare Word Problems. We therefore developed byte-pair tokenization on genomic sequence as a strategy to build multiple frequency balanced vocabularies and select the vocabulary that carries the information content of the human genome in an optimal way. In combination with fine-tuning tasks, and the inbuilt transparency of the model architecture, we can now start using the resulting foundation DNA language model, GROVER (“<u>G</u>enome <u>R</u>ules <u>O</u>btained <u>V</u>ia <u>E</u>xtracted <u>R</u>epresentations”) to extract the different layers of the genetic code with unprecedented performance and detail.

## Results

### Building a frequency-balanced vocabulary on the human genome

Foundation DNA language models require a vocabulary for training that is generated on the genome by grouping nucleotides into tokens. For a human DNA language model this is a difficult task, because sequence composition over the genome is very heterogeneous. While A- and T-rich sequences are relatively frequent, sequences with CG dinucleotides are depleted, due to their susceptibility to mutation ^10^. To achieve tokens of not too heterogeneous frequencies, tokens that contain rarer sequence content should be shorter for optimal contribution to model training, whereas tokens with frequent sequence content should be longer. In the case of CG dinucleotides, this is of particular importance, given that through potential DNA methylation in the form of 5-methyl-cytosine ^11^, this dinucleotide fulfils a special biological role in gene regulation ^12,13^ and retrotransposon silencing ^14^. To tokenize the genome in a frequency-balanced fashion, we employed byte-pair tokenization ^15^, a method originally developed for text compression (**Fig1A**). The algorithm prioritizes building larger tokens of more frequent sequence content by sequentially combining the most frequent token pairs into new tokens. Starting with the four nucleotides A, C, G, and T, in the first cycle of byte-pair tokenization, two Ts are combined to a TT token, where the initial embedding of the token does not retain the information on the sequence content, but adapts a new token identity. This pairing can in principle be continued for many cycles, forming continuously larger tokens. With the dictionary growing, the new pair in every cycle becomes less and less frequent. We use these vocabularies to train a model with BERT architecture (**Fig1B**) for masked token prediction with cross-entropy loss. This results in a model for each cycle of byte-pair tokenization from which the optimal model and therefore vocabulary is selected.

**Figure 1.**
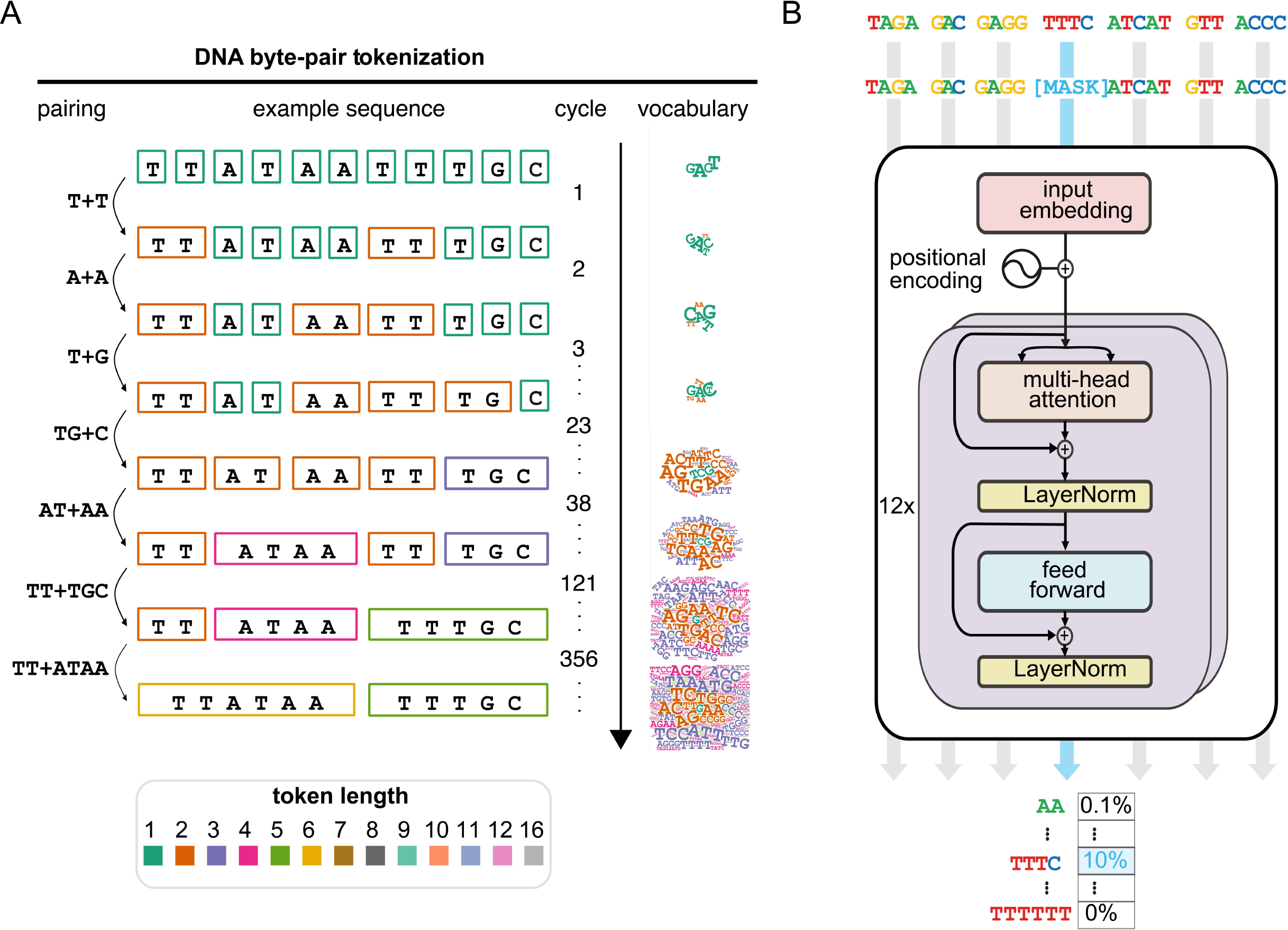
DNA Byte-Pair Tokenization and model architecture. **(A)** The principle of DNA Byte-Pair Tokenization highlighted on an example sequence with the tokenization steps relevant for this sequence. Resulting vocabularies are colored by token length and depicted in a word cloud with relative weights of the words by their frequency. **(B)** The model is a BERT architecture with 12 transformer blocks. The model is embedding the tokens and is trained with cross-entropy loss to predict the masked token and updates the embedding while training. The output is probabilities of the masked token identity.

### Using next-token-prediction to select an optimized vocabulary for DNA language models

To select an optimal vocabulary and model, performance can in principle be assessed with two different strategies, intrinsic validation and extrinsic validation, where extrinsic validation would typically be performed on a task of specific genome biology. Since we aim to build a foundation model that is not biased towards a specific biological task, we consider intrinsic validation more suitable. However, metrics like perplexity and accuracy are dependent on the size of the dictionary and consequently unsuitable for direct comparison of vocabularies. We therefore employed a fine-tuning task of next-k-mer prediction ^9^, which fulfils the criteria of a task that does not directly focus on a specific question of genome biology, is independent of the number of training parameters of the foundation model, the tokenization strategy and vocabulary size. The task is set to predict the next fixed-size k-mer, using 5 fine-tuning models with k=[2,3,4,5,6] and therefore requires knowledge of context for good performance. We applied this task with the foundation models trained on 100 to 5000 cycles of byte-pair tokenized vocabulary and assessed performance through accuracy of next-k-mer prediction (**Fig2A**). While there are small differences between the different k-mer-models, they all generally perform in an optimal range between cycles 400 and 800. We thus picked cycle 600 for our final model, GROVER. To compare GROVER with models of fixed-size vocabularies, we established foundation models of non-overlapping 4mers, 5mers, and 6mers to perform the equivalent task (**Fig 2B**). While they all showed inferior accuracy for next-token prediction, it is noteworthy that foundation models with the matched k-mer size as the next-k-mer prediction task hardly performed better than fine-tuning foundation models with unmatched k-mer sizes. We also assessed performance for the GROVER foundation model training task of masked token prediction (**Fig 2C**), which achieves 21 % accuracy. When allowing for the token to be among the up to 60 top predicted tokens, which represents 10% of the dictionary, accuracy rises to 75%. Perplexity was assessed in comparison to the fixed-size k-mer models, divided by the number of tokens in the dictionary, which to some extent can compensate for the differences in dictionary sizes between the models. GROVER shows perplexity of 12% of the vocabulary, whereas the fixed-size k-mer models show perplexity of 25%, 21%, and 36% for 4mers, 5mers, and 6mers, respectively (**Fig2D**). The selected byte-pair-tokenized vocabulary is therefore outperforming the fixed-size k-mer models and the vocabulary is thus optimized to carry the information content of the genome with relevance for this type of model training. We next had a closer look at the vocabulary composition to ask what characteristics might allow such improvement in performance.

**Figure 2.**
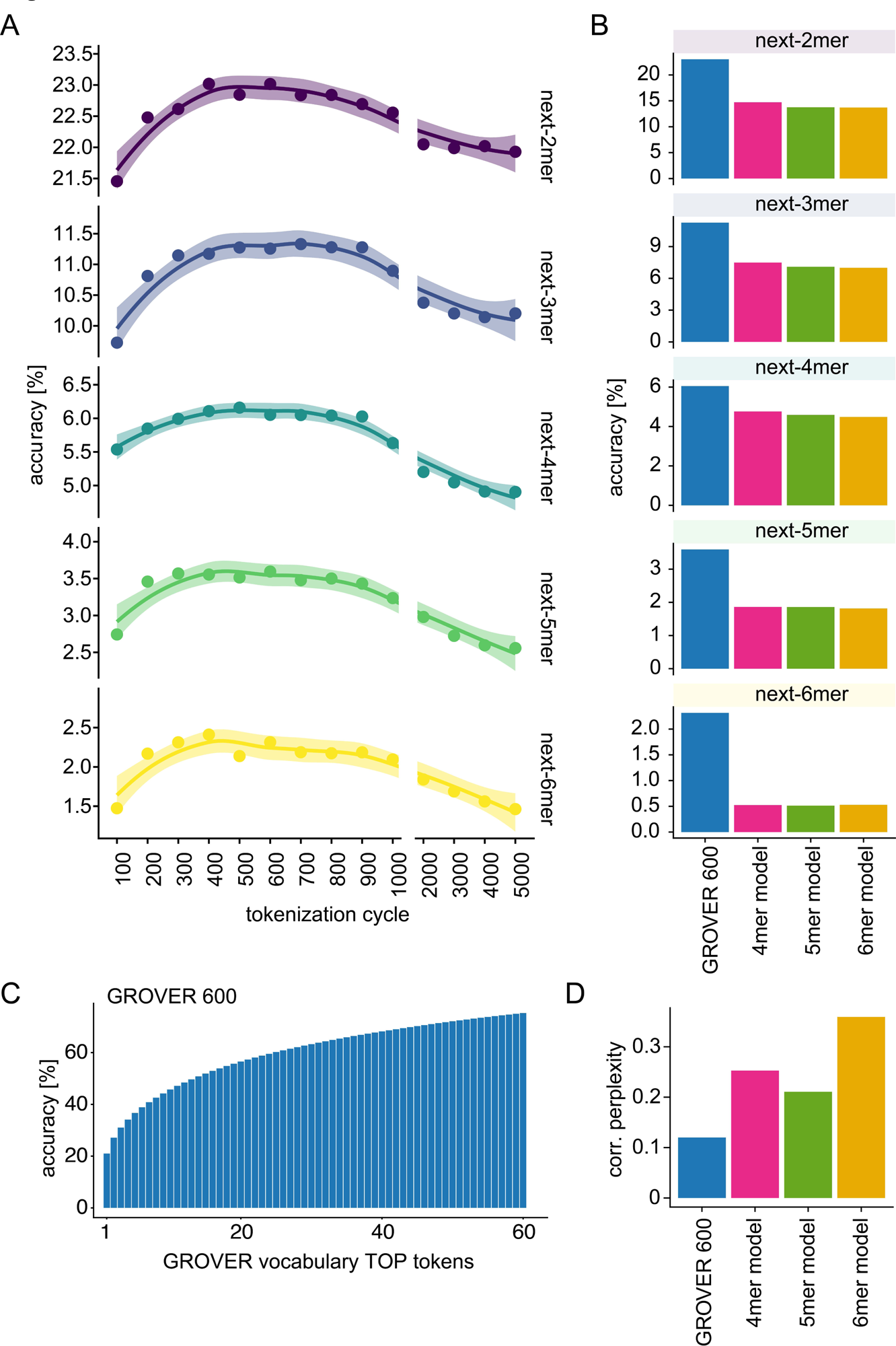
Performance based selection of the vocabulary identifies 600 cycles of Byte-Pair Tokenization as optimal. **(A)** Selection of the optimal vocabulary through accuracy of next-token prediction as a fine-tuning task for the foundation models using prediction of 2 to 6 nucleotide long next-k-mers as readout. Depicted is accuracy with a loess fit and 95% confidence interval. **(B)** Performance comparison using accuracy of next-k-mer prediction as a fine-tuning task. Compared are GROVER with 600 cycles of Byte-Pair Tokenization with models based on k-mer-tokenization, with length of 4, 5, and 6 nucleotides. **(C)** Performance assessment of GROVER with 600 cycles Byte-Pair Tokenization using accuracy for the masked token being predicted as the TOP 1 token, up to TOP 60, i.e. the TOP 10%. **(D)** Performance assessment of GROVER with 600 cycles Byte-Pair Tokenization using perplexity, divided by the total number of words in the dictionary. Comparison with models based on k-mer-tokenization, with length of 4, 5, and 6 nucleotides.

### The vocabulary that carries optimized information content for DNA language training

Byte-pair tokenization started with the special tokens and the four nucleotides. It added one token for each of the 600 cycles. All single As, Cs, and Ts (that were not adjacent to Ns) were incorporated into larger tokens and thus got removed from the dictionary. As a result, the dictionary contains 601 GROVER tokens. First, we assessed their frequency in the human genome (**Fig3A**). While the fixed-size k-mer models show bimodal distributions of token frequencies, the GROVER vocabulary has the vast majority of the tokens with frequencies higher than 100,000 times in the genome with a median of ca. 400,000. While the 4mer model also has most token frequencies above 100,000, the complexity of the dictionary is only 256 tokens as compared to 601 for GROVER. Still, the majority of tokens in the GROVER vocabulary are 4mers and the average token length is 4.07 (**Fig3B**) with a close to symmetric distribution of other token sizes. The length of the tokens ranges from one 1mer, a single guanine, to two 16mers, A_16_ and T_16_. In the formation of the vocabulary not all k-mers are generated, since the required smaller tokens may have been eaten up by other more frequent combinations. This results in a heterogeneous representation of k-mers in the GROVER dictionary (**Fig3C**). Most tokens in the dictionary are 5mers and 6mers with 213 and 224 tokens each and larger tokens become less frequent. The proportional representation of k-mers in the dictionary is therefore also heterogeneous (**Fig3D**). With the G, 25% of the 1mers are represented, and 63% of the 2mers. A CG dinucleotide, for example, has never formed and this sequence is either part of larger tokens or distributed between two tokens. Even though 4mers are the most frequent tokens in the vocabulary, only 32% of the possible 4mer sequences are represented. Still, in total the nucleotide representation within the dictionary is a reflection of the nucleotide composition of the genome (**Fig3E**). There is however a substantial imbalance in respect to the first nucleotide of the tokens in the dictionary with 97% of them starting with an A or a T, while nucleotide composition of the genome would suggest an expected proportion of 60%. This is probably a side-effect of the byte-pair tokenization algorithm, which initially prioritizes the more frequent As and Ts to generate new tokens and thus amplifies the nucleotide frequency imbalance of the genome in the representation of the tokens’ first nucleotides. To characterize the tokens further, we assessed performance metrics per token length and discovered heterogeneous performance. 6mers perform worst on average for the Area Under the Curve (AUC) and accuracy, while both the shorter and especially the longer tokens show much better performance (**Fig3F&G**). Importantly, every token lies above 50% and thus contributes to the non-random performance of GROVER. Interestingly, the tokens with most accurate predictions (>99%), are a 9-mer (ATTACAGGC) and a 12-mer (TGTAATCCCAGC). Least accurate predictions (<1%) are made for a 6-mer (TTTAGG). This heterogenous range of accuracy can either be caused by heterogenous sequence ambiguity of the tokens and their genomic regions, or differences within GORVER’s learning. Therefore, we investigated the relationship between the tokens in the vocabulary and what aspects GROVER learns from the tokens and their context.

**Figure 3.**
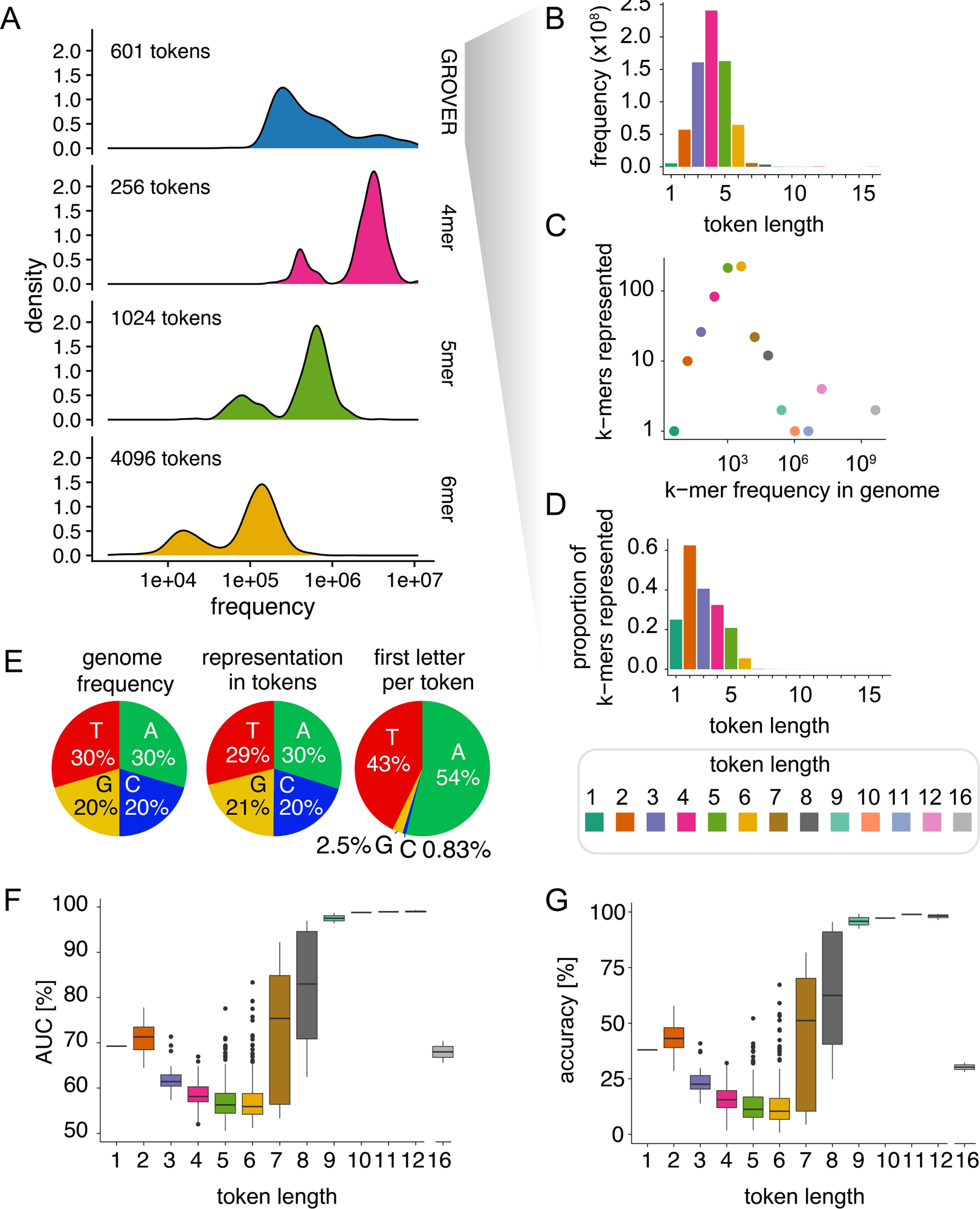
The frequency balanced GROVER vocabulary shows differential learning performance by token length. **(A)** Token frequency in the genome, for GROVER and k-mer-tokenization. **(B)** Composition of the GROVER vocabulary, differentiating token length versus frequency in the genome. **(C)** Composition of the GROVER dictionary, differentiating the frequency of k-mers in the genome and how many k-mers are represented in the GROVER dictionary. **(D)** Composition of the GROVER dictionary, differentiating the proportion of k-mers represented in the dictionary, dependent by length. **(E)** Representation of the four nucleotides in the genome and in the GROVER dictionary. **(F)** Prediction performance as Area Under the Curve (AUC) by token length. **(G)** Prediction performance as accuracy by token length.

### GROVER learns from tokens some of their characteristics and some annotation

From transformer models it can be extracted what the model learns by extracting attention, by analyzing the trained embeddings, and by using specialized fine-tuning tasks. Token embeddings are updated during the training process and thus reflect how the trained model sees the tokens. To interrogate what GROVER learns from the tokens, we took the average embeddings for each token sequence and compared the GROVER embeddings with a Word2Vec embedding ^16^. As a method for static embedding, Word2Vec was built to reflect average word associations in a multidimensional space and thus can be used to assess simple average word associations within the vocabulary. To assess how much context learning can be inferred, we performed a principal component analysis (PCA) and extracted the variance explained from the first principal component (PC1) as a measure of maximum explainable variance (MEV) (**Fig4A**). While Word2Vec embedding shows with 15% a rather high MEV value for an LLM, GROVER lies with 3.5% in a range typical for a model on natural language ^17^. It can thus be concluded that GROVER learns also sequence context and lexical ambiguity that is not reflected in an assessment of average token embeddings. The first two PCs of Word2Vec shows that there are no clear token clusters forming (**Fig4B**). Due to the high variance explained on PC1, its ranks were used to color code the tokens as a visual measure of token similarity, which correlates with GC content (Spearman’s R=0.93) (**Fig4C**) and thus also relates with tokens being present in regulatory elements of the genome, like promoters and enhancers, which tend to have high GC content. Some outlier tokens are also colored blue, despite not being of high GC content. These are predominantly tokens that reside in *Alu* sequences and are thus surrounded by high GC content. A non-linear dimensionality reduction of Word2Vec embedding with Uniform Manifold Approximation and Projections (UMAP) largely reflects the color coding and also does not reveal major token clusters (**Fig4D**). Principal component analysis and UMAP on the trained embedding of GROVER also does not reveal major token clusters. This indicates that GROVER does not simply learn to spell out the tokens ^9^, but the embeddings reflect more complex learned content. The dimensionality reduction reflects the token colors derived from the Word2Vec embedding (**Fig4E & F**) on the second dimension. We therefore asked what the models learn on the first 20 principal components by correlating token characteristics with annotation (**Fig4G & H**). Word2Vec learns GC content on PC1 (Spearman’s R=0.93) and AG content on PC2 (Spearman’s R=0.75). AG content is also a reflection of strand specificity relative to replication, transcription, and direction of SINE and LINE retrotransposons. Although the PCs explain less variance for GROVER, associations with token characteristics and annotation are more pronounced. Since most genome functional annotations are also dependent on GC content, this was corrected for. PC1 strongly correlates with token frequency (Spearman’s R=0.88) (**Fig4I**), which played no visible role for Word2Vec. PC2 correlates with GC content (Spearman’s R=-0.96) (**Fig4J**), similar to Word2Vec’s PC1. Thus, also the colors are reflected. PC3 correlates with AG content (Spearman’s R=0.94) (**Fig4K**) and thus probably reflects the learning of DNA strand information. PC4 and PC6 mildly correlate with token length (Spearman’s R=0.39 and Spearman’s R=0.43, respectively) (**Fig4L&N**). Token length is also related to token frequencies, which may result in the mild correlations. Lastly, PC5 correlates with AC content (Spearman’s R=-0.81) (**Fig4M**), which complements GC and AG content as a measure of token sequence characteristics. Lower principal components show additional mild associations with certain repeat classes, and also gene transcription and replication timing (**Fig4H**). Although GROVER learns with an unsupervised strategy on the human genome, it clearly learns to separate token characteristics through the average trained token embeddings. To interrogate further the token association with general features and annotation with genome function, we performed hierarchical clustering on the Euclidean distance of the average trained token embeddings (**Fig5**). The most prominent cluster is composed on high-frequency 4mers and 5mers. As already indicated with PCA, also GC content, AG content and token length contribute to the distance of the tokens to each other. Assessing the proportion of tokens localizing to repeats shows that there are tokens that almost exclusively do so. They specifically form SINEs, LINEs and simple repeats. While there is also clustering visible for tokens with variable contribution to chromatin colors, beyond repeats there is no clear assignment of tokens to specific genome functional elements. Taken together, GROVER learns from token identity, but learning of functional genomics features needs to rely on a larger sequence context.

**Figure 4.**
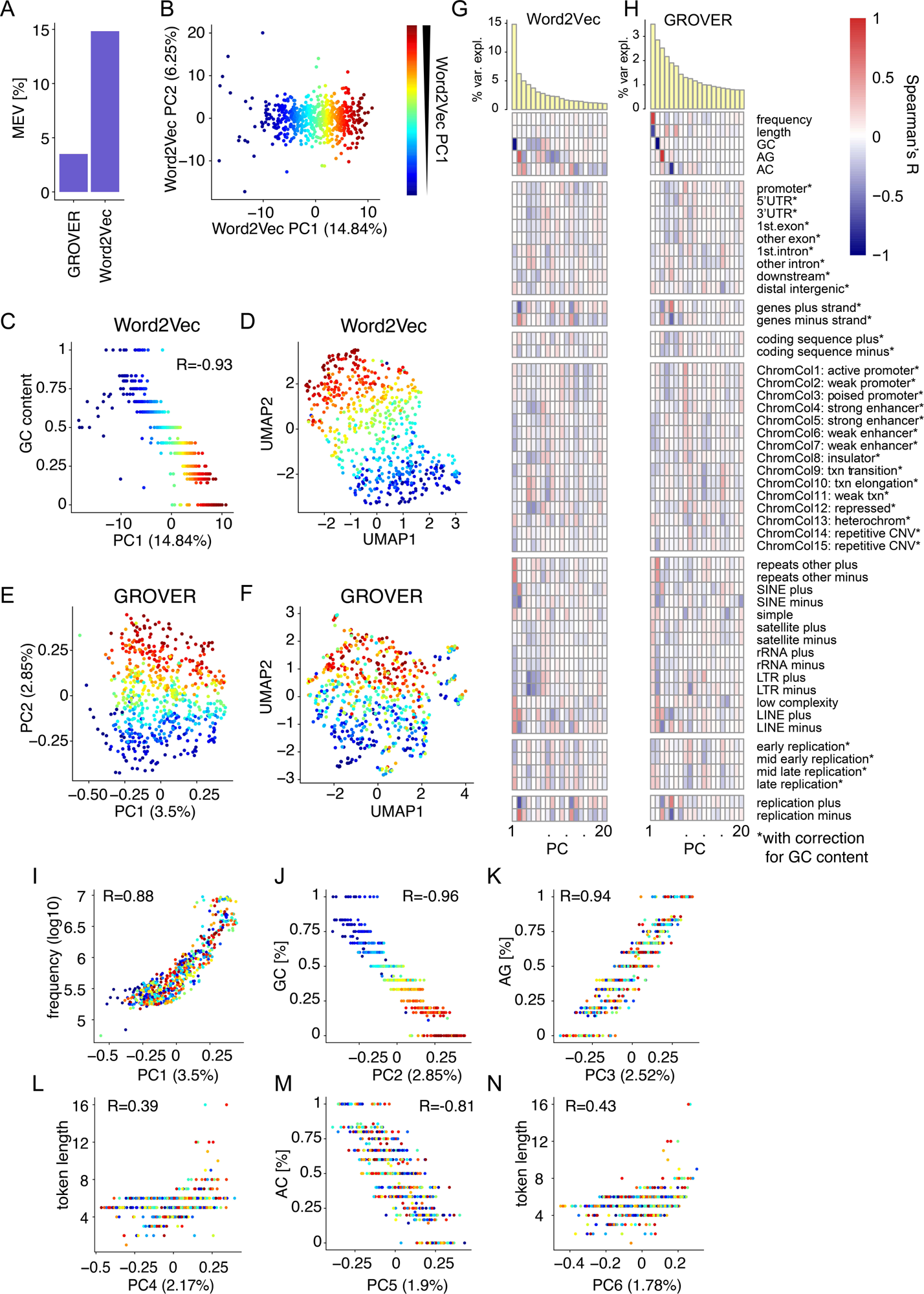
Average GROVER token embedding shows learning of genome information content. **(A)** Maximum Explainable Variance (MEV) derived from the first principal component (PC) of the GROVER embedding averaged for each token, compared to Word2Vec (W2V) static embedding, derived from the same vocabulary. **(B)** Principal component analysis (PCA) of the GROVER vocabulary-derived Word2Vec embedding for the first two principal components with their explained variance. Color represents the rank within Word2Vec PC1. **(C)** Correlation Word2Vec PC1 with token GC content. Color represents the rank within Word2Vec PC1. R = Spearman’s R correlation coefficient. **(D)** Uniform Manifold Approximation and Projections (UMAP) of the Word2Vec embedding of the GROVER vocabulary. Color represents the rank within Word2Vec PC1. **(E)** Principal component analysis (PCA) of the GROVER vocabulary embedding, averaged per token, for the first two principal components with their explained variance. Color represents the rank within Word2Vec PC1. **(F)** Uniform Manifold Approximation and Projections (UMAP) of the GROVER vocabulary embedding, averaged per token. Color represents the rank within Word2Vec PC1. **(G&H)** Correlation of vocabulary characteristics and genome biology annotation of the vocabulary with the GROVER vocabulary-derived Word2Vec embedding (**G**) and with the GROVER token averaged embedding (**H**). Depicted is variance explained (var. expl.) throughout the first 20 principal components (PCs) of a Principal Component Analysis (PCA), along with the Spearman correlation with token characteristics and percentage of tokens of a specific token sequence that belong to genome annotation categories. Gene element annotations with gene promoters (Transcriptional start site +/- 1kb), 5’- and 3’-untranslated regions (UTRs), exons, introns, gene downstream regions (10kb), and distal intergenic regions, as well as gene strand, coding sequence strand, chromatin colors (ChromCol; txn meaning transcribed, CNV meaning Copy Number Variation), replication timing, and replication strand, were corrected for GC content using linear regression. Repeat annotations are obtained from RepeatMasker, SINE=Short Interspersed Nuclear Element, rRNA=ribosomal RNA, LTR=Long Terminal Repeat, LINE=Long Interspersed Nuclear Element. (**I-N**) Correlation of the Principal Components (PCs) from a principal component analysis (PCA) of the GROVER token averaged embedding with features that were identified to explain much of the variance explained in the PC, i.e. PC1 and token frequency (**I**), PC2 and GC content (**J**), PC3 and AG content (**K**), PC4/PC6 and token length (**L** &**N**), and PC5 and AC content (**M**)

**Figure 5.**
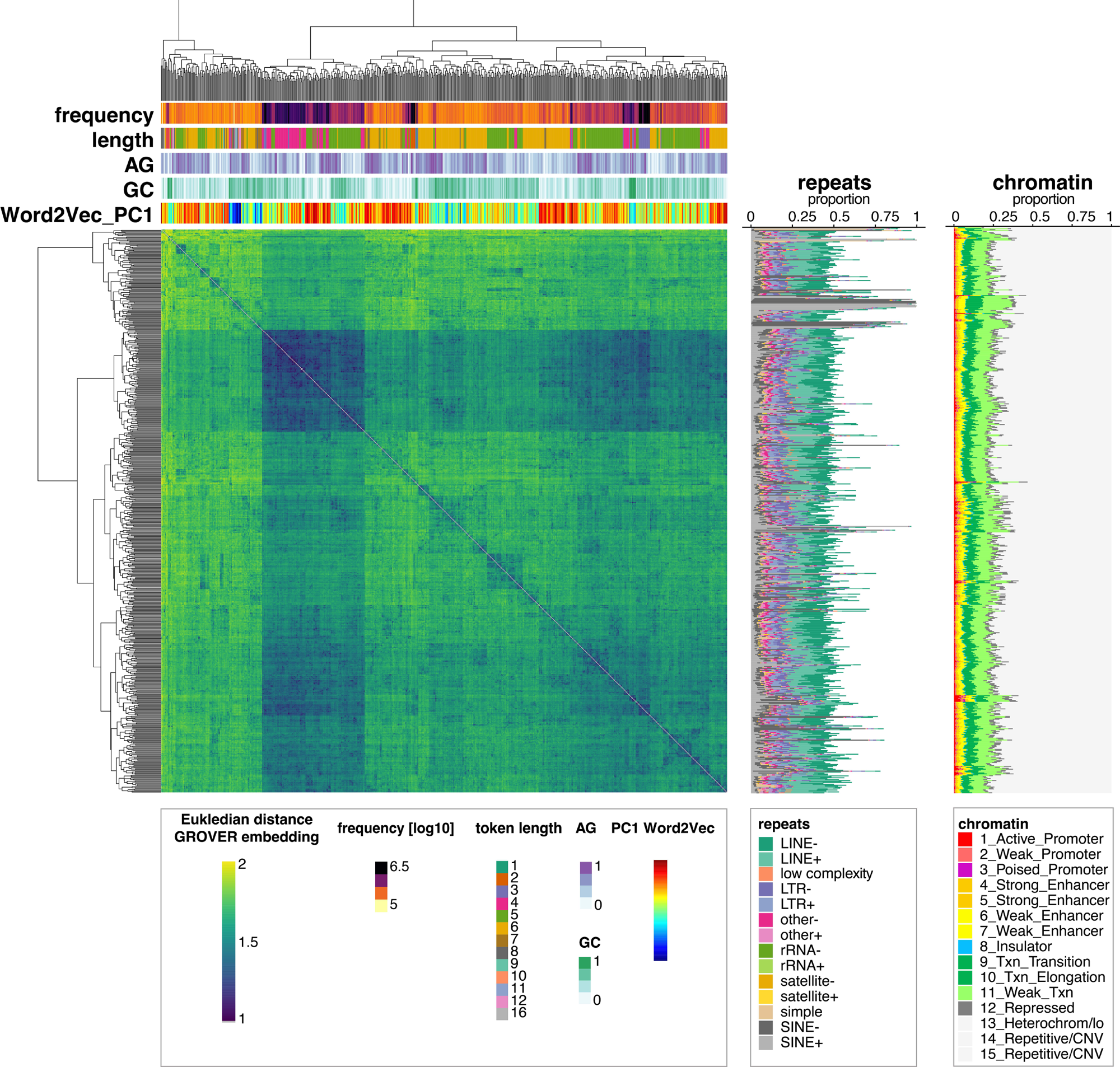
Average GROVER token embeddings cluster by token characteristics and annotation. Average GROVER token embedding, clustered with hierarchical clustering by Euclidean distance. Cluster annotation with token characteristics, i.e. token frequencies in the genome, token length, AG content, GC content, Word2Vec embedding principal component 1 (PC1), proportion of tokens falling into chromatin colors (ChromCol; txn meaning transcribed, CNV meaning Copy Number Variation), and repeats. Repeat annotations are obtained from RepeatMasker, SINE=Short Interspersed Nuclear Element, rRNA=ribosomal RNA, LTR=Long Terminal Repeat, LINE=Long Interspersed Nuclear Element.

### GROVER learns context

We use self-similarity to asses context (**Fig6A**). The self-similarity of a token sequence is the cosine similarity between its embeddings across its different contexts. The more contextualized the representations are, the lower we expect self-similarity. Using hierarchical clustering on the Euclidean distance over self-similarities for each transformer layer, clustering resembles the analogous results of the average embeddings. Self-similarity varies between the layers. Tokens that localize largely to repeats show highest self-similarity almost throughout the layers. They are also distinct in their good per-token performance metrics and long token lengths. Otherwise, high self-similarity in layer 12 is also highlighting a special token group, the short tokens of high frequency, which also show good performance metrics. For the other tokens it can be concluded that dependent on their cluster, there is low self-similarity in different layers, which indicates contextualized learning. To investigate whether there is sufficient context that reflects particular genome biology, we used the average embedding per window of 510 tokens over the genome to interrogate which regions GROVER identifies as similar or distant. The windows correspond to an average sequence length of 2.1 kb (**Fig6B&C**). Non-linear dimensionality reduction with UMAP shows that there is a spread of sequence context with some little clusters but no clear separation of larger groups. Annotation of repeats over the windows (**Fig6B**) indeed shows that some of the smaller clusters can be explained by presence of particular repeats, such as a cluster that associates with LINE elements in antisense to the tokenization direction, or a small cluster with association to satellite repeats. In general, however, most repeats are not particularly enriched in any area of the UMAP 2D space. Annotation to chromatin colors (**Fig6C**), however, shows distinct locations for some chromatin features in the larger cloud, such as the different types of promoters and repetitive sequences. Within these features there has not been a strong enrichment of particular token sequences (**Fig6A**), which suggests that annotation of the windows in the genome have been learned through context. This shows that GROVER can learn biological information directly from the sequence and suggests its suitability for further fine-tuning tasks to address genome biology.

**Figure 6.**
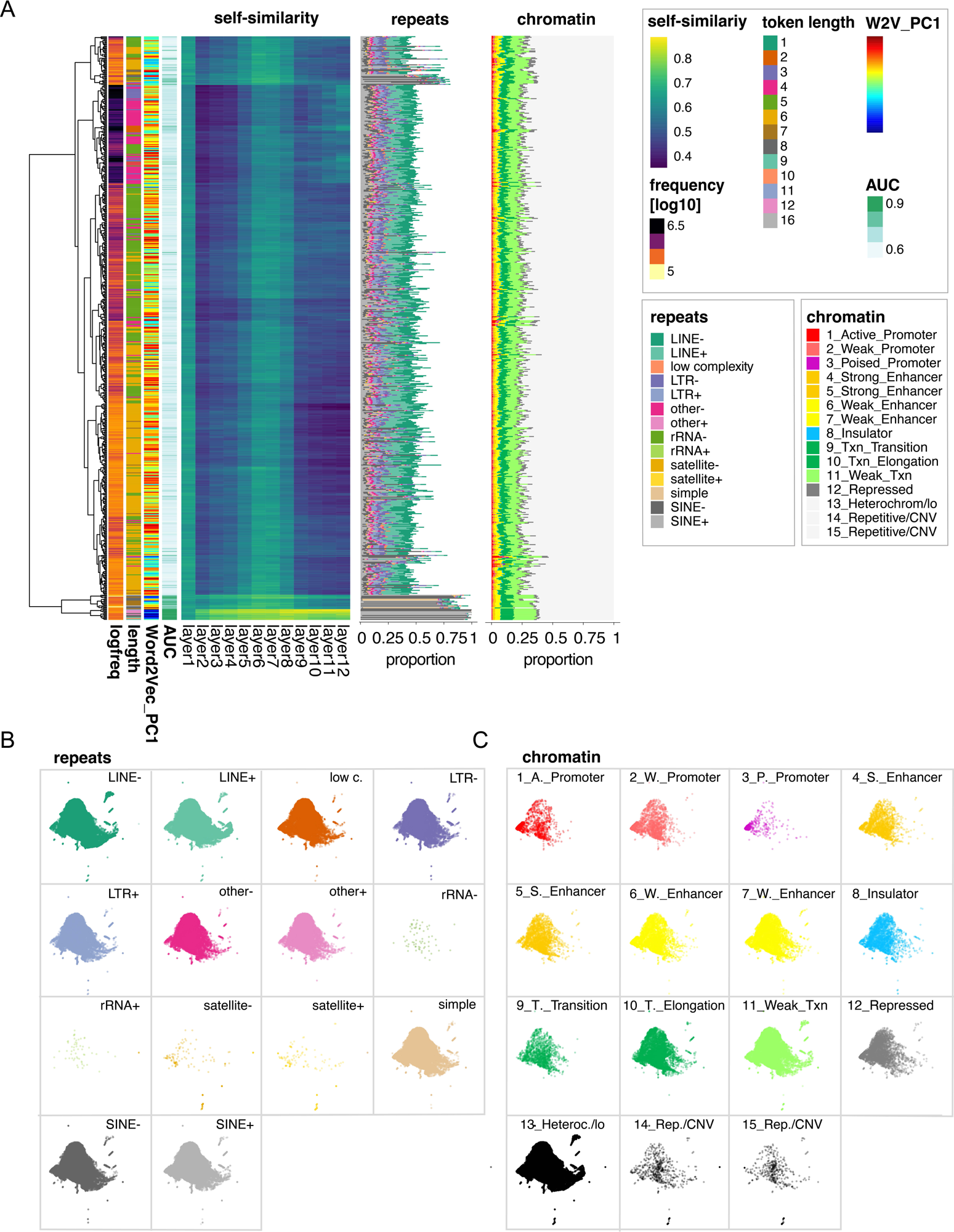
GROVER learns token context and genome annotation. **(A)** Self-similarity per token sequence as extracted by cosine similarity of the same token in different contexts throughout the 12 transformer layers. Self-similarity is clustered with hierarchical clustering by Euclidean distance. Cluster annotation with token characteristics, i.e. token frequencies in the genome, token length, performance (area under the curve; AUC), Word2Vec (W2V) embedding principal component 1 (PC1), proportion of tokens falling into chromatin colors of ChromHMM (ChromCol; txn meaning transcribed, CNV meaning Copy Number Variation), and repeats. Repeat annotations are obtained from RepeatMasker, SINE=Short Interspersed Nuclear Element, rRNA=ribosomal RNA, LTR=Long Terminal Repeat, LINE=Long Interspersed Nuclear Element. **(B&C)** Uniform Manifold Approximation and Projections (UMAP) for the average embedding of regions in the genome 510 tokens in size. Regions are annotated relative to repeats (**B**) and chromatin colors (**C**), and shown, if they overlap with the feature of interest.

### GROVER can be fine-tuned for tasks to understand genome biology

To show the suitability of GROVER for genome biological questions, we selected three representative fine-tuning tasks. Prom-300 was adapted with some minor modifications from Ji et al.^6^ (**Fig7A**). In short, promoters are selected for sequences around transcription start sites (TSS) -250 bp/+50 bp and classified as actual promoters versus promoters with shuffled tokens. For shuffling relative to the GROVER tokens, the task performs with an F1 score of 0.996 as compared to 0.90, 0.89, and 0.81 for the k-mer models of 4mers, 5mers, and 6mers, respectively, and 0.83, 0.74, and 0.69, when using their own k-mers for token shuffling. (**Fig7B**). A more challenging task is Prom-scan (**Fig7C**), where 1 kb windows are selected from the 10 kb regions around the TSS and classified for overlap with the TSS. Due to unbalanced classes, this task is more challenging, yet GROVER recognizes the TSS windows with an F1 score of 0.60 as compared to 0.51, 0.49, and 0.45 for the k-mer models (**Fig7D**). Lastly, we approached a task of protein-DNA binding focusing on CCCTC-Binding Factor (CTCF) (**Fig7E**).

**Figure 7.**
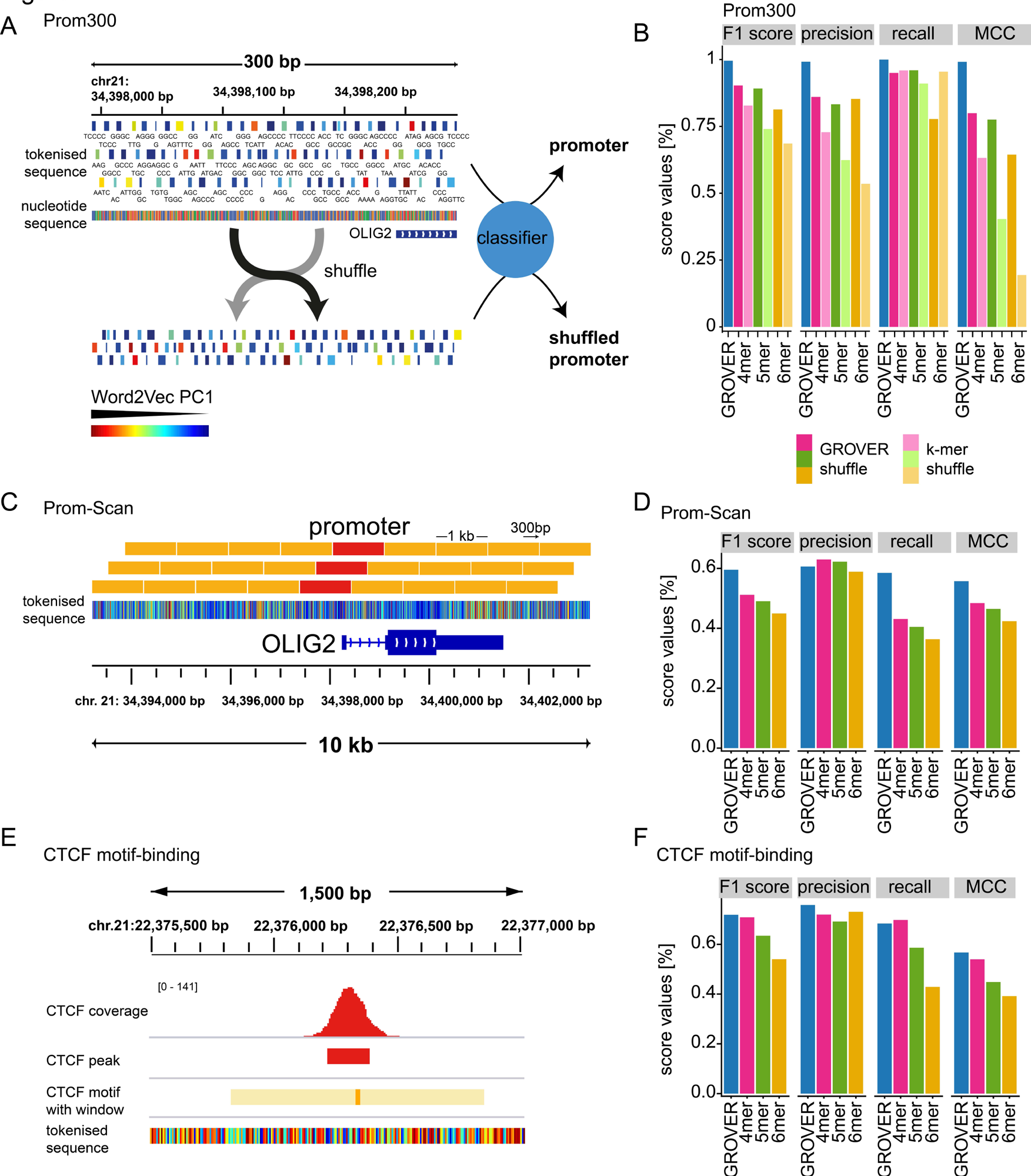
GROVER outperforms models with fixed-size k-mers for biological fine-tuning tasks. **(A)** Prom300 promoter classification, where promoters are defined as sequences around transcriptional start sites (TSS) with -50 and + 250 base pairs. As negative class, tokens are shuffled in 6 randomly selected chunks of 8 equally sized chunks within promoters. As a fine-tuning task the promoters are classified into real promoters versus shuffled promoters. The *OLIG2* promoter is shown in the context of the nucleotide sequence with A in green, C in blue, G in yellow and T in red. GROVER tokens are colored by their rank in principal component 1 (PC1) of Word2Vec embedding, which explains ca. 15% of the variance within a static embedding of the vocabulary and thus can serve as a visual distance measure of the tokens, which correlates strongly with GC content. **(B)** Performance for the Prom300 task of GROVER versus models of fixed-size k-mers with the shuffling based on GROVER tokenization and tokenization that matches the respective benchmarking models. Metrics are given as F1 score, precision, recall, and Mathews Correlation Coefficient (MCC). **(C)** For Prom-Scan promoter assignment, windows are defined in regions of 1001 bp around the TSS +/- 10kb with an offset of 300 bp. Within these 1001 bp windows, the fine-tuning task classifies overlap with the TSS. Shown is the *OLIG2* promoter with GROVER tokenization, colored by their rank in principal component 1 (PC1) of Word2Vec embedding. **(D)** Performance of the Prom-scan task of GROVER versus models of fixed-size k-mers. Metrics are given as F1 score, precision, recall, and Mathews Correlation Coefficient (MCC). **(E)** CTCF-binding prediction task, where regions are defined as CTCF-binding motifs +/- 500 bp and GROVER is trained for recognizing whether such motifs are actually bound by CTCF, determined via ChIP-seq peak-calling in HepG2 cells. Depicted is a random region in the genome that has a prominent CTCF binding peak overlapping with a CTCF binding motif, in the context of GROVER tokenization, colored by their rank in principal component 1 (PC1) of Word2Vec embedding. **(F)** Performance of the CTCF-motif-binding task of GROVER versus models of fixed-size k-mers. Metrics are given as F1 score, precision, recall, and Mathews Correlation Coefficient (MCC).

The task is to recognize which sites that contain a CTCF binding motif are indeed bound by the protein using ChIP-seq data from HepG2 cells obtained from ENCODE ^18^. While there are ∼85,000 binding motifs in the human genome, only ∼32,000 are indeed bound by CTCF. Beyond the motif, sequence context with particular physicochemical properties of the DNA, as well as binding of other proteins, determines whether a protein indeed binds its assigned motif. GROVER achieves for this task an F1 score of 0.72 as compared to 0.71, 0.63, and 0.54 for the k-mer models (**Fig7F**). This shows that interestingly the 4mer model is getting closest in performance and also generally tends to outperform the 5mers and 6mers.

GROVER does not only achieve good performance with tasks that are agnostic to genome biology questions, but also achieves good performance with tasks that directly address genome function, particularly those, where sequence context needs to be learned. It is still an open question how protein-DNA binding is generally encoded in the DNA beyond direct binding motifs. The larger context of how protein binding is encoded in the genome as well as other research questions of genome function can now be addressed through extracting the learned representations from GROVER.

## Discussion

We have built GROVER (“<u>G</u>enome <u>R</u>ules <u>O</u>btained <u>V</u>ia <u>E</u>xtracted <u>R</u>epresentations”), a foundation DNA language model with an optimized vocabulary for the human genome, selected by a task of next-k-mer prediction, a fine-tuning task that is independent on the structure of the foundation model and thus can handle different vocabulary sizes and tokenization strategies, without directly selecting models for biological tasks. GROVER can grasp DNA language structure by learning both characteristics of the tokens and larger sequence contexts. It outperforms analogous models with fixed-size k-mers both for tasks of next token prediction and fine-tuning tasks of promoter identification and DNA-Protein binding. Thus, we have identified the vocabulary that currently best defines the information of the human genome as it can be extracted by a BERT model. GROVER can be a basis to extract the information content of the genome, its grammatical and general language structures via token embeddings or through extracting attention from the foundation model. In addition, token embeddings and attention can be obtained from specific fine-tuning tasks to interrogate specific genome biology. Such tasks could be genome annotation with functional data, genotype-phenotype predictions, or technical tasks for example for data augmentation.

We have built GROVER exclusively on the human genome, which is a different strategy from other models, such as Nucleotide Transformer ^7^, where different genomes are combined in one model. While the number of parameters is limited by using just one genome, the models serve different purposes given that different genomes follow different language rules. For example, the human genome contains *Alu* retrotransposons to ca. 12%, which are primate specific genome elements. While the coding genome is relatively conserved over multiple species, the non-coding genome is more unique. It is these language rules we aim to learn with GROVER in a transparent way, specifically for biomedical questions and therefore decided not to mix the human genome with other species. Still, also with this strategy we are not compromising in regards to performance for biological fine-tuning tasks, where a one-species approach seems to be sufficient. However, this approach may hit its limits with smaller genomes.

The different layers of the genetic code can now be approached through these models and it can be extracted how DNA is coding for protein and transcripts, for genes regulation, self-propagation, and stability. In there lies not only the key of genotype to phenotype prediction and the information of what in the DNA makes us human, but also the information of predisposition to disease and treatment responses, which is to a large extent encoded in the patients’ general and somatic genomes. DNA language models like GROVER therefore have the potential to substantially push progress in personalized medicine.

## Methods

Unless otherwise specified as being written in R, all code is written in Python.

The final source code and fine-tuned models will be made available at https://zenodo.org

### Data

We used the *Homo sapiens* (human) genome assembly GRCh37 (*hg19*) and only take into account the sequences that contain A,C,G and T. Each chromosome is split into windows varying between 20 and 510 tokens in size. Specifically, with a 50% probability, the size of a window is 510. With another 50% probability, its size is a random integer between 20 and 510. 80% of the windows are taken as training set and the remaining as test set.

### Byte Pair Tokenization

Byte Pair Encoding (BPE) or Byte Pair Tokenization has originally been developed as a text compression algorithm ^15^ and is now widely adapted as a tokenization strategy for transformer models on natural language like GPT-3 ^4^. We adapted the tokenizer from Sennrich et al ^19^ for genome sequence and used up to 5000 cycles of tokenization.

Tokens are visualized in a Word cloud with the R package Wordcloud2 (https://github.com/Lchiffon/wordcloud2).

The optimal vocabulary was selected through performance assessment with next-k-mer prediction with GROVER

### The DNA Language Model GROVER

GROVER adapts the Transformer Encoder BERT architecture ^8^. It takes as input tokenized sequences of up to 510 tokens long. In addition to the vocabulary generated from the genome, GROVER takes five special tokens: CLS, PAD, UNK, SEP and MASK. These special tokens are commonly used in language models, CLS represents the classification token, PAD is used for padding the right side of the sequence in case it is shorter than the maximum input length of the model, UNK is for sequences of nucleotides that do not belong to the vocabulary, SEP is used to indicate the end of a sequence and MASK represents the masked tokens. The model is trained as a masked language model. In the implementation aspect, masking a token means replacing the original token for the MASK token. Conceptually, the MASK token hides this part of the sequence to the model.

In a given input sequence, 2.2% of the tokens are selected, of which 80% are substituted with a special mask [MASK] token, 10% of tokens are randomly replaced with standard tokens (i.e. any token different from the class [CLS], pad [PAD] or mask [MASK] token).

To pretrain the model, we gather more than 5 million samples from the genome. Training was carried out on clusters of A100 GPUs, on batches of sizes 64 with an Adam optimizer and a learning rate of 4^-4, epsilon 10^-6, beta 0,99, maximum input length of 50, dropout probability of 0.5, batch_size 64, and 0.022 probability of masking.

### Next k-mer prediction

For model validation and selection of the optimal vocabulary we used a fine-tuning task of next-k-mer-prediction that we previously developed ^9^. It allows to compare different foundation models that rely on context learning independent on how their vocabulary was generated, the size of the vocabulary, or the learning parameters. The task is not dependent on a specific biological question. The principle of next-k-mer-prediction is to take the pre-trained language models and fine tune it to predict the next k-mer, where k is 2, 3, 4, 5 and 6.

Chromosome 21 is split into sequences of 510 nucleotides, where we keep the first 56 nucleotides of each sequence. We randomly select 500,000 sequences, 80% for training and 20% for testing.

The samples are defined as the first 50 nucleotides of each sequence. For the labels, we take the k nucleotides that follow the 50 nucleotides. The next-kmer model has 4^k different classes, i.e., 16, 64, 256, 1024 and 4096, respectively, which are all the permutations of k nucleotides. We use an Adam optimizer with learning rate 10^-6, epsilon 10^-8, beta 0,99, maximum input length of 50, dropout probability of 0.5, and batch_size 64.

From the models we extract performance metrics and use accuracy for token prediction for the decision of the optimal vocabulary and comparison to other tokenization strategies. For comparison of Byte-Pair Tokenization cycles and visualization with ggplot2 (3.4.1) in R (4.2.1) we use a loess fit with a 95% confidence interval.

### kmer models

For the k-mer models, we use the same parameters and samples as GROVER, only different by the vocabulary and tokenization. Tokenization is performed with not-overlapping k-mers, 4, 5 and 6 nucleotides in length. The vocabularies consist of all the permutations of k consecutive nucleotides (i.e. 256, 1024 and 4096 respectively) as well as the five special tokens described above.

### Word2Vec

For comparison of token embeddings, we use Word2Vec a static word embedding tool ^16^ that maps each word to a single vector. In general, this mapping function does not account for lexical ambiguity, that identical letter sequences can have multiple interpretations or different grammatical roles. We use Word2vec with a continuous bag-of-words (CBOW) approach for learning representations of words. CBOW predicts the middle word from surrounding words in a sentence with a small number of words as context. The order of the words is not taken into consideration. To generate the Word2Vec (W2V) embeddings, 300,000 sequences are randomly chosen from the training set and use the Word2Vec module of Gensim (https://radimrehurek.com/gensim/models/word2vec.html), with the following parameters: min_count = 1, vector_size = 768, window = 5.

### Model Embedding

We obtain a contextualized word representation that is the token embedding of the BERT model. To obtain the token embeddings of the model, we extract the weights of the layer word_embeddings for each kmer model. Where needed, we derive either the embedding of all 12 transformer layers, a summarized version for each token sequence, or a summarized version for sequences of 510 token lengths.

### Dimensionality reduction

We obtain dimensionality reduction from average token embeddings that are represented as vectors with length 768. Principal Component Analysis (PCA) and Uniform Manifold Approximation and Projections (UMAP) were performed in R (4.2.1) with the packages ‘stats’ (4.2.1) and ‘UMAP’ (0.2.10.0), respectively with default parameters. Maximum Explainable Variance (MEV) as a measure for context learning ^17^ was extracted as the variance explained by the first principal component. Clustering of the token embeddings and self-similarity was performed with hierarchical clustering of Eucledian distance with pheatmap (1.0.12).

Dimensionality reduction for extracted embeddings of genome windows was performed in Python. 100,000 bins of 510 tokens were randomly selected and the 768 dimensions of the embedding matrix were reduced to 50 dimensions by PCA and further reduced to two dimensions using UMAP. UMAP was configured with standard parameters, including 15 nearest neighbors and a minimum distance of 0.1.

### Self-similarity

Self-similarity was assessed as the cosine similarity of different embeddings from the same token sequence, separately for all 12 transformer layers of the BERT architecture.

5,000 embeddings per token were gathered from the test set and pairwise cosine similarity for each token was computed in every layer.

### Genome Annotation

Tokens and sequence windows were annotated to token characteristics and functional genomics data in R (4.2.1) with the GenomicRanges (1.50.2) package, for genome information BSgenome.Hsapiens.UCSC.hg19 (1.4.3) and TxDb.Hsapiens.UCSC.hg19.knownGene (3.2.2). Sequence was derived with Biostrings (2.66.0) to obtain GC, AC, AG and nucleotide content, as well as k-mer frequencies. Gene element annotation was performed with ChIPSeeker (1.34.1). Regression of GC content was performed using the residuals of a loess regression from the stat (4.2.1) package. Strand annotation relative to transcription was obtained from TxDb.Hsapiens.UCSC.hg19.knownGene (3.2.2). ChromHMM annotation was used from the GM12878 lymphoblastoid cell line from ENCODE, downloaded via the UCSC genome browser (https://www.genome.ucsc.edu/), where also the repeat masker was obtained for annotating repeats. Replication strand and timing were obtained from OK-Seq data from K562 chronic myeloid leukemia cells from the reanalysis by Pongor *et al.* ^20^ downloaded via GEO (GSE131417). Tokens were annotated by determining the proportion of tokens overlapping with a specific annotation. Genome regions were annotated by determining overlaps.

### Fine-tuning task promoter identification, Prom300

Promoter sequences were obtained from the EPD database (https://epd.epfl.ch/), and lifted over to the hg19 assembly. Promoters were defined as the Transcriptional Start Sice (TSS) +249/-50 bp. The corresponding Byte Pair sequences were retrieved from the Byte-Pair-tokenized genome or converted into k-mer tokenized sequence with k=[4, 5, 6]. For generating negative class samples, each tokenized sequence was split into 8 chunks, and the tokens of six randomly selected chunks were shuffled among those six chunks. For the classification task for correct or shuffled sequence, the dataset was split 80%-10%-10% for training, validation, and test.

### Fine-tuning task promoter scanning, Prom-Scan

The same human promoter annotations as used in the Prom300 task were taken in 10 kb windows around the TSS. The sequence was divided into overlapping 1001 bp windows with a step size of 300 bp. Training was performed for classification of the presence of a TSS in a respective window. Only the central TSS was considered, even in the presence of more than one TSS. The dataset was split 80%-10%-10% for training, validation, and test.

### CTCF motif binding

CTCF ChIPseq peaks in HepG2 cells were derived from the ENCODE project (https://www.encodeproject.org/experiments/ENCSR000BIE/). The human CTCF motif was retrieved from the JASPAR database (https://jaspar.genereg.net/matrix/MA0139.1/), and significant motif occurrences in the hg19 genome derived with FIMO from the MEME suite. Motifs that intersect with a CTCF peak were considered to be CTCF bound. Classification was performed for bound and unbound CTCF motifs with a split 80%-10%-10% for training, validation, and test.

### Data and code availability

The code for paper reproduction: https://doi.org/10.5281/zenodo.8373203

A tutorial on how to use GROVER as a foundation model: https://doi.org/10.5281/zenodo.8373159

Pretrained GROVER: https://doi.org/10.5281/zenodo.8373117 The tokenised hg19 genome (600 cycles): https://doi.org/10.5281/zenodo.8373053

### Author contributions

ARP has conceptualized the study, MS and JH applied the models and implemented the fine-tuning tasks. All authors designed the fine-tuning tasks and analyzed the data. ARP wrote the manuscript with input from MS and JH.

## Acknowledgement

This work was supported by the Center for Scalable data analytics and artificial intelligence (Scads.AI) Dresden-Leipzig. **ARP** was supported by the Mildred Scheel Early Career Center Dresden P2, funded by the German Cancer Aid. **MS** was supported by a TU Dresden Junior Fellowship and a Maria Reiche Postdoctoral fellowship of TU Dresden. **JH** was supported by the TU Dresden program ‘FOSTER – Funds for Student Research’.

